# The influence of larval retention on coral recruitment

**DOI:** 10.1101/2025.03.14.643210

**Authors:** Marine Gouezo, Peter Harrison, George Roff, Aaron Chai, Damian Thomson, Magda Guglielmo, Lauren Hardiman, Alicia Forbes, Brint Gardner, Christopher Doropoulos

**Affiliations:** Faculty of Science and Engineering, Southern Cross University, Military Road, East Lismore, 2480, NSW, Australia; CSIRO Environment and CSIRO IMT Scientific Computing Services, St Lucia, Queensland, Australia

**Keywords:** slack current, reef-scale hydrodynamics, coral, larvae, coral recovery, supply, settlement, tiles, reef substrata, density-dependence

## Abstract

Marine broadcast spawners typically exhibit bipartite life-histories with distinct pelagic larvae and benthic phases. The transition between phases shapes benthic populations, but the rate of larval arrival to a reef is largely unknown due to challenges in accurately measuring supply. Once larvae arrive to a reef, reduced current flow and velocity, facilitate the transition from the water column to the benthos for inefficient swimming larvae. Yet, for coral reefs characterised by complex hydrodynamics and tides, slack current conditions typically last 1.5-3 hours and it remains unclear if such short retention periods drive significant recruitment. This study mechanistically examined the effects of water retention on the settlement of coral larvae from the water column to the benthos and subsequent longer-term recruitment over 15-months. Brief periods of slack currents (<3-hours) retained larvae in unconstrained larval supply treatments, resulting in settlement rates 40-times higher than natural, background rates. Constrained and longer retention of larvae under nets for 2.5- and 24-hours resulted in 4-7-times higher initial settlement than the unconstrained treatment and 305-times higher than background rates. However, after 15-months, similar numbers of surviving recruits were observed across all larval supply treatments, highlighting the effects of density-dependent population regulation. Observations from recruitment tiles show survival rates of coral recruits after 15-months were low (<0.25), even though gregarious settlement behaviour and settlement close to tile edges improved survival. In contrast, observations from the natural substrate show survival rates were 2.5-3.5-times higher than tiles after 15-months, indicating density-independent survival due to optimal niche space and less space limitation. Therefore, when larval supply is high and gregarious behaviour prominent, key vital rates including recruitment and mortality derived from settlement tiles are likely overestimated, as substrate and microhabitat properties between tiles and natural reef environment vary. Overall, our study highlights the prominent role of slack current conditions and local retention of larvae in facilitating the supply-to-settlement transition, and how this interacts with density-dependent processes post-settlement. Our findings underscore the need to investigate how the interaction strengths of pre-and post-settlement processes modulate early coral recovery to best model recovery trajectories for conservation and restoration prioritisation.

## INTRODUCTION

Larval supply is a critical process influencing population dynamics of benthic marine invertebrates. Most produce pelagic larvae that disperse in the water column before reaching competency and settling on suitable reef substrate (Harrison and Wallace 1990, Bertness et al. 1999). The rate of larval arrival to a reef is the first process shaping population dynamics and has long been a key focus in benthic ecology (Roughgarden et al. 1988, Caley et al. 1996, Hughes et al. 2000). The temporal dynamics of larval arrival to reefs, also referred to as larval fluxes, are known to be influenced by parental spawner biomass, propagule biological characteristics, larval predation, and hydrodynamic processes, all of which are key pre-settlement factors (Jones et al. 2009, Pineda et al. 2010) (Fig. 1a). The magnitude and frequency of larval fluxes to a reef have predominantly been quantified in temperate ecosystems (e.g. Minchinton and Scheibling 1991, Olivier et al. 2000), and only sporadically in tropical systems such as coral reefs (Black 1988, Oliver et al. 1992, Gilmour et al. 2009), limiting our understanding of how larval supply operates in coral reefs (Fig. 1a).

**Figure 1.**
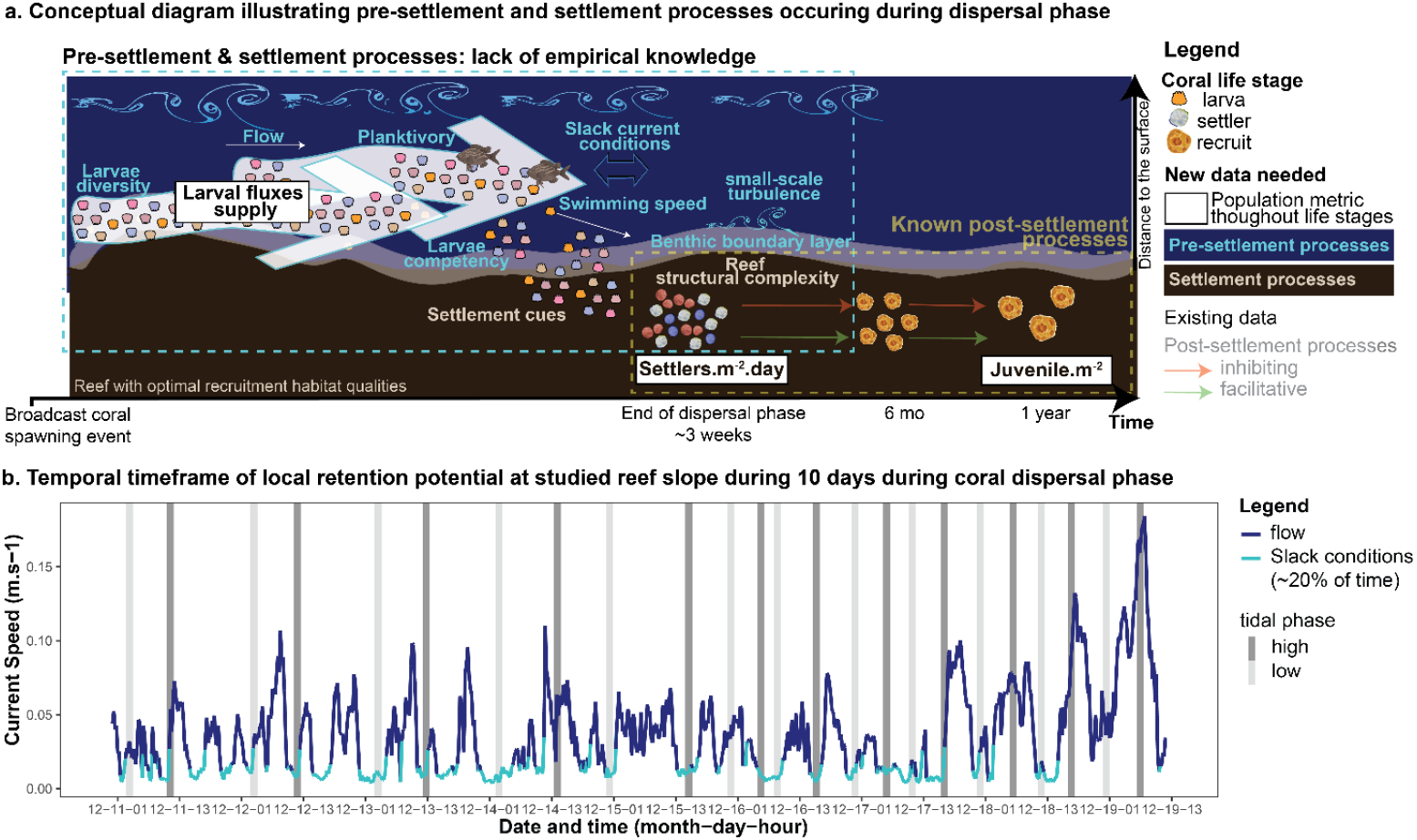
(a) Conceptual diagram emphasizing pre-settlement processes where empirical data are currently lacking and (b) current speed profile at the study reef showing timeframes of slack current conditions (current speed <0.015 m.s^-1^) occurring ∼20% of the time.

For coral reefs, there is a clear lack of empirical data on processes influencing propagule arrival to a reef during their planktonic phase and during their transition to the benthos (Pineda et al. 2010; Fig. 1a). Most research on larval supply from broadcast spawning corals has relied mainly on dispersal modelling, with few studies quantifying larval supply *in situ* throughout the dispersal phase (Black 1988, Oliver et al. 1992, Gilmour et al. 2009). Reef-scale hydrodynamics are inferred through hydrodynamic modelling, *in situ* current measurements, or a combination of both, and are challenging to characterize as they occur from large to small scales (e.g. Wolanski and King 1990, Bruyère et al. 2023). From this knowledge, we understand that the spatio-temporal patterns of water movement on a reef following the arrival of a larval plume can lead to localized retention (Wolanski et al. 2024, Gouezo et al. 2025). Bays, gulfs and inlets sometimes retain these plumes for >24 hours that leads to high self-recruitment rates (Golbuu et al. 2016, Wolanski et al. 2024). However, most coral reefs do not have the hydrodynamic characteristics of bays and inlets. Instead, they experience intermittent periods of fast and slack current conditions; the latter typically lasting 1-3 hours on average (Fig. 1b), leading to episodic conditions of local retention (Gouezo et al. 2025). During periods of rapid water movement, coral larvae may experience difficulty swimming towards the reef due to their inefficient swimming ability (Hata et al. 2017). Conversely, when currents become slack (Fig. 1b), larvae have more opportunities to transition from the pelagic environment to the benthos and search for settlement substrate on the reef. However, it remains unclear if short slack current periods are sufficient to drive demographically relevant coral larval settlement. When propagules are close to the benthos, they may become entrained in the benthic boundary layer where flow speeds generally decrease, but can be heterogenous due to microturbulence, fine-scale eddies, and interactions with benthic structures (Fig. 1a) (Boudreau and Jorgensen 2001). In addition, the ability of propagules to transition from the water column to settle on the benthos are driven by their biology and ecological interactions, processes that are partly understood through experimental laboratory and field studies (e.g. Reidenbach et al. 2007, Connolly and Baird 2010) and then extrapolated into modelling studies investigating reef recovery trajectories (e.g. Bozec et al. 2022, Lachs et al. 2024).

From settlement to one year of age, biophysical interactions influencing coral recruitment success are well studied (Fig. 1a; (e,g, Doropoulos et al. 2016, dela Cruz and Harrison 2017, Gouezo et al. 2020). Many corals selectively settle on or near particular, facilitative substrata (Harrington et al. 2004, Arnold et al. 2010) and in cryptic microhabitats that provide refugia from predation by from fish and invertebrates (Babcock and Mundy 1996, Penin et al. 2011). Larvae can exhibit gregarious settlement to enhance fusion among related individuals (Grosberg and Quinn 1986), leading to chimerism that increases growth and survival (e.g. Raymundo and Maypa 2004, Puill-Stephan et al. 2012; dela Cruz and Harrison 2017). However, high settler density can intensify competition (Vermeij and Sandin 2008, Doropoulos et al. 2017) and attract predators, increasing relative mortality rates (Gaines and Roughgarden 1985). This may trigger a negative density-dependent response, which stabilizes populations at carrying capacity (Hixon and Johnson 2009). Abiotic drivers also influence ecological trade-offs during recruitment (Doropoulos et al. 2016), with sedimentation, microhabitat complexity, currents and light intensity having prominent roles in driving patterns of recruitment (Harriott and Fisk 1987, Babcock and Davies 1991, Maida et al. 1994).

Processes regulating the earliest phases of coral populations from larvae to one-year-old juveniles have primarily been investigated using settlement tiles because newly settled corals are tiny (<1mm) and typically settle cryptically, requiring stereomicroscopy for quantification (dela Cruz and Harrison 2017; Edmunds 2023). However, artificial substrates inadequately represent natural reef microhabitats and benthic communities due to biases related to materials, structural complexity, orientation, and conditioning time (Harper et al. 2021, Hamizan et al. 2021). Coral recruitment quantified using tiles are primarily used as proxies (Edmunds 2023) but often parametrised into ecological models and extrapolated to ecosystem processes (e.g. Bozec et al. 2022, Lachs et al. 2024). Elucidating patterns of coral recruitment on natural reef substrata is needed to understand population dynamics during earliest life-stages but challenging due to the microscale and cryptic nature of recruitment (Doropoulos et al. 2016, Harrison et al. 2021; Edmunds 2023). Macrophotogrammetry provides a tool to assess key vital rates of recruitment, growth, and post-settlement survival on natural reef substrata for all coral genera (Gouezo et al. 2023). Understanding vital rates from larval to juvenile stages is particularly important as this cryptic phase drives recovery (Caley et al. 1996, Doropoulos et al. 2022) and is a focal point in recovery-based management and coral reef restoration (Gouezo et al. 2021, Banaszak et al. 2023).

In this study, we explored the effect of water retention and larval density on larval settlement, growth, and survival. We manipulated retention timeframes and densities of coral larvae, and monitored coral settlement and post-settlement survival and growth on tiles and natural reef substrate for 15-months. Through three experiments, we tested whether (1) typical slack current conditions on reefs sufficiently retain larvae to contribute to localized settlement and longer-term recruitment, and (2) how larval density influences immediate settlement and longer-term recruitment. We then applied macrophotogrammetry monitoring techniques over >12-months following supply to (3) directly assess coral recruitment on the natural reef substrate to determine whether vital rates measured using settlement tiles accurately reflect benthos occurrences. Finally, we (4) examined factors affecting coral recruitment on tiles to understand differences in recruitment rates between tiles and reef, focusing on population processes (chimerism and density-dependence) and abiotic characteristics (spatial location and light exposure).

## MATERIALS AND METHODS

### 1. Study location and coral larvae cultures

The study was conducted on a fringing reef habitat along the northwest flank of Lizard Island (−14.650070°, 145.450139°), a mid-shelf reef in the northern Great Barrier Reef. In 2022, coral spawn slicks were collected from the *Acropora*-dominated reef slope between Palfrey and South Island, within a few hours after spawning on December 10, 2022. Gametes and embryos were gathered using pool scoops with 150 µm plankton mesh and transferred to a 250-l collection tub onboard (Harrison 2024). Gametes and embryos were then placed into a 4 × 4 m larval pool with a 3 × 3 m net for culturing (Harrison 2024), anchored in the sheltered south-eastern lagoon of Lizard Island in ∼10 m water depth. Coral larvae were cultured for five days until >50% were competent. On December 15, 5.5-day-old larvae were concentrated and transferred into a 250-l container. Five x 15 ml samples were taken for larval density estimates. The concentrated larvae culture was divided into equal volumes of ∼50,000 larvae per sample. Samples were placed in heavy duty food-grade plastic bags (5-l) filled with medical grade oxygen, and stored in shaded 32-l containers for 1-hour before transport to the experimental site.

### 2. Experimental design

Twelve experimental plots (1.5 m x 1.5 m) were established on the reef slope, randomly positioned on either side of a 50-m transect at 5-m depth following the reef’s contour (Fig. S1a). Each plot was marked with galvanized deck nails at both ends of its diagonal. Four larval retention treatments were randomly assigned to the 12 plots, with 3 replicates per treatment. Three control plots were established ∼300 m upstream. A tilt current meter (Lowell TCM-4) was installed near the transect midpoint to monitor currents for ∼10 days during the experiment. One day before larval releases, 5 × 5 × 1 cm preconditioned travertine limestone settlement tiles were prepared (Fig. S1). Each tile was assembled on a stainless-steel bolt with a bird band secured underneath. Stainless-steel baseplates with welded nuts were nailed to the substrate (Fig. S1) along the plot diagonals, then tiles were secured onto the baseplates. After deployment (6-8 per plot), 1.5 m x 1.5 m net enclosures of 150 µm plankton mesh were set up over the 12 plots and secured using tent pegs and chain links.

Each enclosure featured a zippered top section (Fig. S1, Harrison et al. (2021)). Prior to larval releases, all enclosures were left open. On December 15, ∼20 min before adding larval retention treatments, all enclosures were closed. Competent larvae were supplied to each experimental plot within 30 minutes during slack current conditions (Fig. S2), following the design for Experiments 1 and 2 (Fig. S3).

*Experiment 1* investigated the effect of retention timeframes on coral settlement and longer-term recruitment. 100,000 larvae (∼44,000 larvae per m^2^) were supplied to each of the 9 plots: 3 x open plots (current = 0.01 m s^-1^ for 2.5-h), 3 × 2.5-h netted plots, and 3 × 24-h netted plots. For the open plots without nets, larvae were dispersed directly over the plot, ∼30 cm above the substrate. Following larval supply, divers carefully left the area to minimize local turbulence near the reef. After 2.5- and 24-h, dives were conducted to zip-open netted plots, which were left on site.

*Experiment 2* investigated the effect of larval density on coral settlement and longer-term recruitment. The experiment supplied two larval density treatments within the 2.5-h retention scenario: 50,000 larvae (∼22,000 larvae per m^2^) and 100,000 larvae (∼44,000 larvae per m^2^) per plot. Three replicate plots were utilised per treatment following the same procedure as in Experiment 1. Control netted plots with no larvae supplied were closed after the larval retention treatments were established and were opened 2.5-hours later (Fig. S3).

### 3. Coral settlement and recruitment monitoring

After 48-hours, settlement tiles were collected from each plot and placed in a basket to prevent damage to early-stage settlers. A stereomicroscope was used to examine and count coral settlers on all tile surfaces. The top and bottom faces of each tile were photographed using a Sony αriv camera with a LAOWA 24mm f/14 2X Macro Probe lens. The following day, tiles were returned to their plots for ongoing monitoring. Subsequent observations were conducted at 66-, 255-, and 455-days post-larval supply. During monitoring, all tile surfaces were inspected under a stereomicroscope and high-resolution images were captured of the top and bottom faces. For the 66-day monitoring, the same camera setup was used as for the initial settlement. For the 255-and 455-day monitoring, once recruits were larger, an Olympus TG6 (12MP) camera in an Olympus PT-059 housing with a 3000 lumens LED ring light (Kraken Weefine) mounted around the objective was used and photos were taken *in situ*.

Image analysis tracked the location and size of each settler over time, differentiating between experimentally seeded and natural settlers and assessing chimerism rates. A batch-processing Python workflow cropped, edited, and labelled tile photos across time points. Visible recruits were annotated using the polygon tool in CVAT (Sekachev et al. 2023). Resulting annotation files were processed using Python (see codes provided via dropbox link), where each recruit polygon was classified as alive, dead, natural recruit, or chimera based on its location within the tile grid over time. Chimerism was determined when more than one recruit from a monitoring time point was found within the perimeter of a single recruit at the subsequent monitoring time point.

Visual monitoring of coral recruitment on reef substrates within a year after larval supply events is difficult due to the small size and cryptic settlement of most recruits (Roth and Knowlton 2009). To address the objective of Experiment 3, macrophotogrammetry was applied in two of the larval retention treatments to quantify coral recruitment on reef substrates, following Gouezo et al. (2023). Survey efforts focused on the longest larval retention treatments (2.5-and 24h-net, 44,000 larvae m-2) as continuous sampling for all treatments was logistically unfeasible. Macrophotography captured the same reef substrate location, covering an average area of approximately 570 cm2. The equipment included a Sony αriv camera with a Sony 90 mm FE Macro G OSS lens, in SeaFrogs housing with a 67 mm threaded flat port for a 90 mm macro lens. To enhance illumination, an LED ring light was secured around the housing port, and four 2000 lumens underwater video lights (Scubalamp PV22) were mounted on a PVC quadrat using flexible arms. Images were batch-edited using Affinity photo software (Affinity Serif LTD, 2021). Following the macrophotogrammetry workflow described in Gouezo et al. (2023), 3D reconstructions were processed using Agisoft Metashape (Agisoft LLC 2021) on the CSIRO High-Performance Computer. Visible coral recruits were annotated on the 3D models by analysing high resolution imagery, their 2D areas measured, and compared over time to assess survival and growth. The approach was also conducted at 2-3 days post-settlement; however, newly settled corals with minimal calcification structure could not be accurately identified on high-resolution images. Therefore, data from 66-days and onwards are presented in this study.

### 4. Data analysis

As recruitment was primarily dominated by acroporids due their dominance within the supplied larval assemblages, the following analyses were conducted excluding the few data on merulinids (n = 2), pocilloporids (n = 8) and others recruits (n = 2).

#### 4.1. Larval retention effects on coral recruitment (Objective 1)

To investigate the effect of larval supply retention treatments on coral settlement, the count of 2-3 day old settlers on tiles was modelled as a function of treatments (4 levels: control, open plot, 2.5-h retention, 24-h retention) using a generalized linear mixed effect model (GLMM) with negative binomial distribution, including survey plots as a random term. Control tiles had zero settlers from 66-days onwards so were removed from subsequent analysis, which modelled recruit count as a function of treatments (3 levels: open plot, 2.5-h retention, 24-h retention), time post-settlement in days (4 levels: 2, 66, 255, 455) and their interaction using the same GLMM to explore the effect of seeding treatment through time.

To test if larval supply retention treatments influenced recruit survivorship on tiles over time, recruit status (alive, dead) was modelled as a function of seeding treatments for each time period (2-66 d, 66-255 d, 255-455 d) using a GLMM with binomial distribution (Bernoulli), including tiles and survey plot as random terms. Average proportional survival per time period and cumulative proportional survival (tile level) were plotted to visualize findings.

#### 4.2. Larval density effects on coral recruitment (Objective 2)

For experiment 2, the same analytical approach was conducted where we investigated the effect of larval density (3 levels: 0 larvae m^-2^, 22,000 larvae.m^-2^, 44,000 larvae.m^-2^) within the retention timeframe of 2.5-h on the count and survival of coral recruits on tiles.

#### 4.3. Substrate effects on coral recruitment (Objective 3)

Recruitment, post-settlement survival, and growth were quantified on tiles and natural reef substrate within two larval retention treatments (2.5- and 24-hour retention, 44,000 larvae.m-2). We tested for significant differences in recruit counts between treatments using a LMM with treatment and time as fixed effects and sample area (log-transformed) as an offset. Since no significant effect of larval supply retention on coral recruit counts was found on either substrata (Table S7) or tiles (Table S1), samples were pooled for subsequent analyses.

Recruit counts were modelled as a function of substrate type, time, and their interaction, with sampled area (log-transformed) as an offset variable, and survey plots as a random effect, using a GLMM with negative binomial distribution. The sampled area on substrata ranged from 150 to 1130 cm^2^, which was 3 to 16 times larger than the tile area (70 cm^2^).

Consequently, more recruits were expected in larger sampled areas. Log-transforming the sampled area ensured larger areas led to a proportional increase in recruit counts. To visualize results, recruit numbers were standardized to m^2^.

Recruit survival (alive, dead) was modelled as a function of substrate type (2 levels: tile, reef) for each period (66-255 days, 255-455 days), using a GLMM with binomial distribution, including survey plots as a random effect. Proportional survival was plotted to visualize findings.

Fusion of coral recruits was only detected on tiles. Recruit growth rate (mm^2^.mo^-1^) was evaluated for two intervals: 66 to 255 days, and 255 to 455 days. Growth rates were analysed based on substrate type and fusion occurrence, with categories: absence of chimeras (reef), presence of chimeras (tiles), and absence of chimeras (tiles). A Gamma-log family distribution GLMM was used, with survey plot as a random effect. Some recruits showed negative growth rates, contradicting Gamma-log family assumptions. The likelihood of shrinkage and its frequency in tile versus reef environments were examined. With no significant difference and shrinkage probabilities of 0.05 (66-255 days) and 0.2 (255-455 days), the growth model was applied to positive growth data points only (removing 13 negative data points) to satisfy Gamma-log family assumptions.

#### 4.4. Tile-specific drivers of mortality (Objective 4)

Recruit survivorship (alive or dead) between 2 and 66 days was modelled as a function of initial settler density (2 days post settlement). Between 66 and 255 days, and 255 and 455 days, it was modelled as a function of recruit density recorded at the beginning of each period, chimerism (present or absent), and recruits’ distance from tile edge. Interactions between chimerism with density and with distance were initially examined but found to be non-significant, so the model was simplified to additive. Analyses used GLMMs with binomial distribution, with tile replicate and survey plots as random effects.

Given that chimerism was identified as a significant factor in survival between 66 and 255 days, we conducted further analysis on how initial settler density and larval retention treatments affected chimera numbers on tiles. We used a GLMM with Poisson distribution to examine the relationship between chimera count per tile with recruit density at 2 days and supply treatments, at the tile replicate level. We applied the same model using recruit density at 66 days, as chimera formation occurs later (Puill-Stephan et al. 2012). To account for the non-linear relationship between chimera counts and recruit density, we incorporated natural cubic splines into the Poisson GLMM when modelling recruit density.

All statistical analyses were conducted in R version 4.3.2 (R Development Core Team 2023). GLMM analyses were done using the glmmTMB package (Magnusson et al. 2017).

Dispersion and model residuals were checked and validated using the package DHARMa (Hartig 2017).

## RESULTS

### 1. Larval retention effects on coral recruitment on tiles

Retention treatments had a highly significant effect on initial larval settlement on tiles (GLMM, P<0.001) (Fig. 2, Table S1-2). Settler density was 40 to 305 times higher across all larval retention treatments (8 - 61 settlers per tile on average) than control tiles (0.2 settlers per tile on average). Settlement was 4 to 7 times higher in the 2.5-h (43 settlers per tile) and 24-h (61 settlers per tile) retention treatments, respectively, compared to the open treatment (8 settlers per tile). While settlement was highest in the 24-h retention treatment, it was not significantly different from the 2.5-h retention treatment (Table S1).

**Figure 2.**
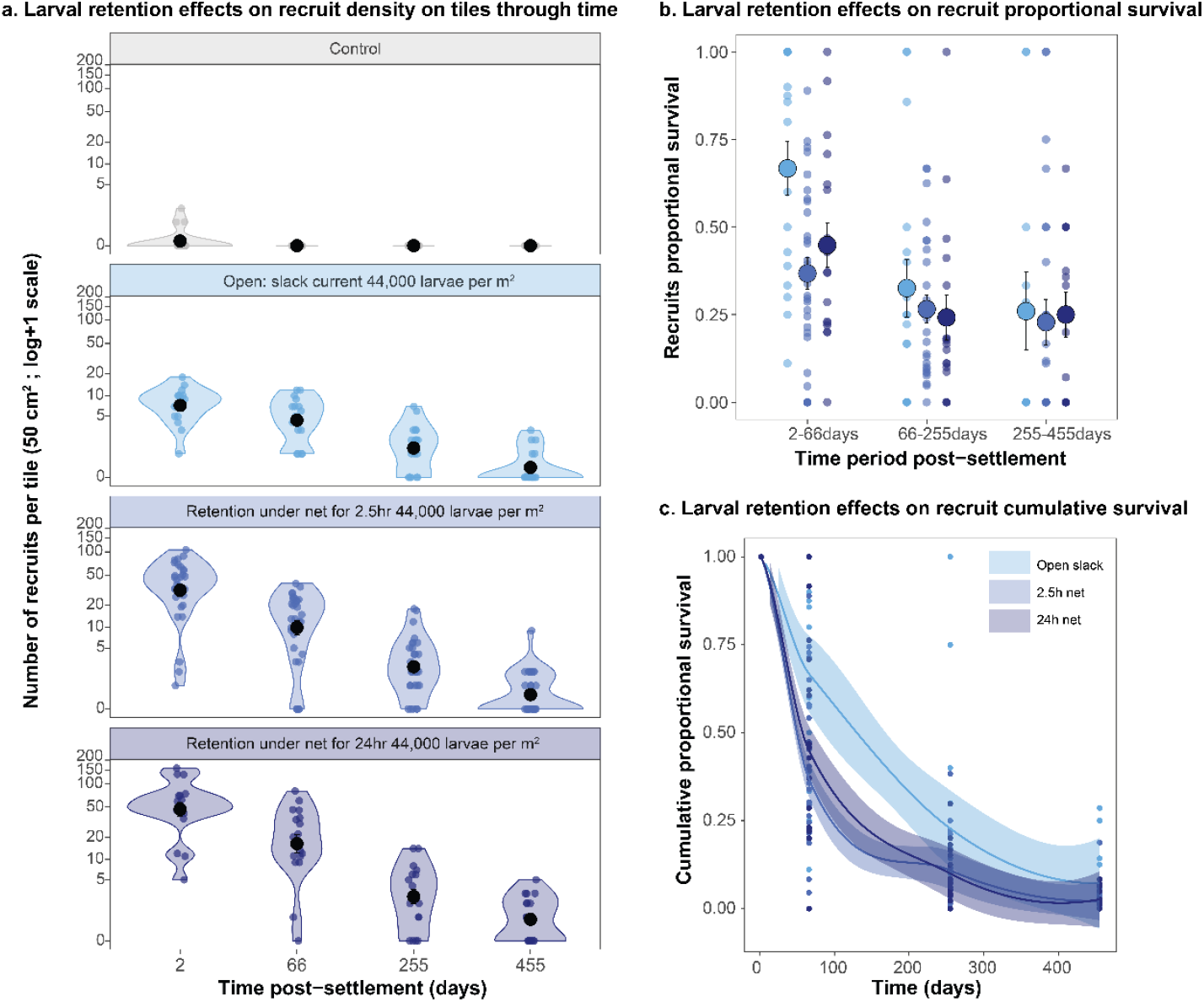
Larval retention treatments effect on coral recruit (a) densities, (b) proportional survival, and (c) cumulative survival on tiles.

From 66 days onwards following settlement, while the number of recruits was always higher than controls (i.e., zeros), no significant differences were observed among retention treatments (Fig. 2). There was a marginally significant effect of retention treatment, with recruit counts higher in the 24-h retention than the open treatment (GLMM-NB, P=0.07, Table S2). Recruit density on tiles decreased significantly over time (GLMM-NB, P<0.001, Table S2), from an average of 5-25 recruits per tile after 66 days to 0.5-1.3 recruits per tile after 455 days.

From 2-66 days, the survival rates of early-stage recruits on tiles varied significantly across seeding treatments (GLMM, P<0.05, Table S3). Recruits from the open treatment averaged 0.66 survival, 1.5-1.8 times higher than those under net treatments averaging 0.4 survival (Fig. 2, Table S3). Following, there were no differences in survival rates among treatments from 66-255 days or 255-455 days post-settlement (Table S3).

### 2. Larval density effects on coral recruitment on tiles

The effect of the two larval density treatments (22,000 and 44,000 larvae per m^2^) within the 2.5-hour retention period on settler density on tiles was significantly greater than the control group (GLMM-NB, P<0.001, Fig. S4, Table S4). However, no significant difference was observed between the two larval density levels (Table S4), which averaged ∼38 spat per tile.

From 66 days following settlement, while the number of recruits per tile was always higher than the controls (i.e., zeros), the number of recruits in the two density treatments remained similar (GLMM-NB, P>0.05, Table S5). Recruit density decreased over time (GLMM-NB, P<0.001, Table S5), from an average of 12-15 recruits per tile after 66 days to 0.5-0.9 recruits per tile after 455 days (Table S5). Concurrently, survival of post-settlement recruits on tiles did not differ between the two larval densities through time (GLMM, P>0.4, Table S6).

### 3. Contrasting coral recruitment, survivorship and growth rates on the reef and tiles

Recruitment rates to the reef did not differ between the two focal larval retention treatments (2.5h-net and 24h-net; Table S7) but were significantly lower than on tiles (Fig. 3a, Table S7-8, Fig. S5). The difference was 11-fold after 66 days, 5-fold after 255 days, and 3-fold after 455 days, implying space-dependent density-dependent limitation on the tiles compared to the space-independent reef substrate (Fig. S5). At 455-days post-settlement, the recruitment rate on the reef was 47 (±7.6) individuals m^-2^, compared to 162 (±34.8) individuals m^-2^ on the tiles. A significant interaction between substrate type and time was observed for coral recruit counts. While recruit numbers on tiles decreased significantly over time, the reef showed similar recruit counts at 255 and 455 days (Table S8). Recruit survival was significantly lower on the tiles than the reef (GLMM-binomial, P<0.01, Table S9), averaging ∼0.25 for the 66-255- and 255-455-days periods for recruits on tiles compared to 0.86 and 0.62 for the recruits on the reef, respectively (Fig. 3b).

**Figure 3.**
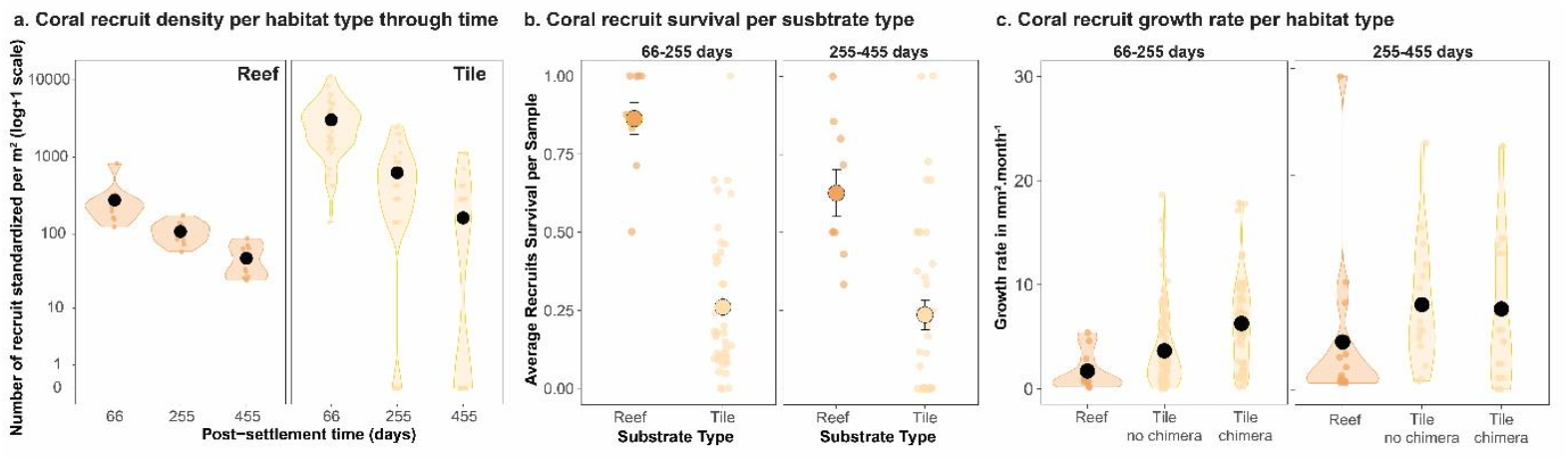
Acroporidae coral density m^-2^ (a) survival (b) and growth rate (c) between the reef and tile substrate.

In the first 255 days, coral recruits on the tiles demonstrated 2.2-to 3.8-fold higher growth rates to those on the reef (Fig. 3c, Table S10). However, from 255 to 455 days, growth rates were similar across both substrate types, averaging 4.5 to 7.9 mm^2^ per month (Fig. 3c). Fused coral recruits observed exclusively on tiles exhibited 1.7-times faster initial growth rates than non-fused recruits in the first 255-days after settlement (Table S10).

### 4. Tile-specific drivers of coral recruit mortality

The survival of recruits during the period from 66 to 255 days was significantly impacted by both chimerism and their location on the tile underside (GLMM-Binomial P < 0.05, Table S11, S12, Fig. S6a). Chimeras demonstrated almost 100% survival; 1.7-fold higher than non-fused recruits. Moreover, a negative relationship between recruit survival and distance from the tile edge was observed, averaging ∼0.7 at the edge and declining to ∼0.5 near the centre (Fig S7a). In contrast to chimerism and settler location, recruit densities did not have a significant influence on survival of recruits on the tiles from 66-255 days (GLMM-Binomial, P > 0.05, Table S12). However, a negative density effect pattern can be seen (Fig. S7) but is likely masked due to many tiles with zero survivors. For the subsequent period between 255 and 455 days, no factors showed significant influence on recruit survival (Table S11, S12).

The number of chimeras on the tile was positively influenced by the initial number of settlers at both 2- and 66-days post-settlement (Table S13, Fig. S6b). The relationship was non-linear, showing a peak at 50 settlers at 2 days and at 25 recruits at 66 days. While the number of chimeras appeared higher in the net retention treatments, this finding was not statistically significant (Fig. S6b).

## DISCUSSION

Once larval plumes arrive on a reef, pre-settlement mechanisms occurring at fine spatio-temporal scales can facilitate the supply-to-settlement transition. These mechanisms include interactions among reef-scale hydrodynamics, topography, propagule biological and behaviour characteristics, benthic complexity, and settlement cues from the reef (Fig. 1). Our study examined whether slack current conditions on reefs, which typically last 1.5-3 hours, can temporarily retain larvae locally, resulting in elevated settlement and subsequent recruitment. Our findings demonstrate that episodic slack currents effectively retain some larvae locally and increase coral larval settlement rates (open treatment) at higher rates than low supply controls, thereby facilitating supply-to-settlement transitions. When larvae were fully retained under nets for 2.5- or 24-hours, settlement densities were 4-7-times higher when compared to the unconstrained releases. However, after 15-months, recruit densities were similar across all larval retention treatments due to density-dependent limitation occurring to settlers from the 2.5- and 24-hour larval retention treatments during the initial 2-months (Fig. 4). In space limited and cryptic microhabitats on settlement tiles, gregarious settlement and settlement close to the tile edges improved post-settlement survival (Fig. 4b). Regardless, on the reef substrate where space limitation was relaxed and recruitment niches optimal, survival rates were 2.5- to 3.5-times higher compared to the tiles, demonstrating how density-dependent processes interact with early population survival in specific microhabitats (Fig. 4a, 4b).

**Figure 4.**
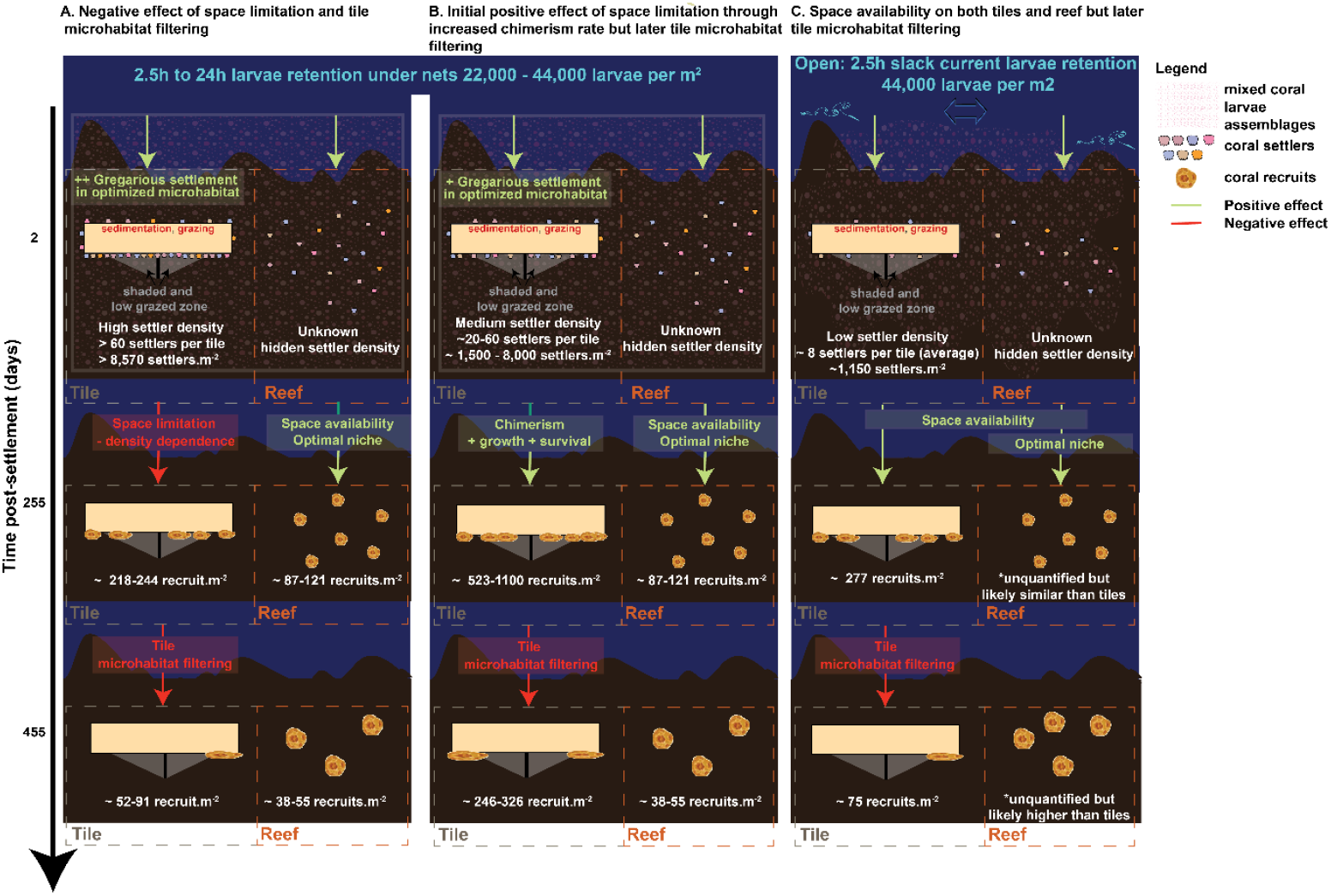
Diagram summarizing the study findings

Our study also addressed a long assumed but previously undemonstrated phenomena: artificial tile substrates inadequately represent natural reef substrates (Harper et al. 2021, Edmunds 2023) and were found in this study to overestimate initial settlement rates on reef substrata due to their optimized initial settlement microhabitats. This pattern was especially apparent when larval retention was constrained under nets for longer periods (i.e., 24-h) and gregarious behaviour of coral larvae led to hypersaturation (Fig. 2, S4). As a result, tiles failed to provide an accurate representation of recruitment on reef substrates in this study, which possess a great diversity of microhabitats (Roth and Knowlton 2009, Doropoulos et al. 2016). Therefore, when the reef provides optimal microhabitats for settlement and survival, the ratio of surviving recruits per competent larvae is much higher than settlement tiles (Fig. 3) as space limitation is relaxed, niche space is optimal, and density-dependent regulation more limited. This highlights how vital rates derived from settlement tiles may be inaccurate and skew the assessment of coral colonization dynamics, which are fundamental in understanding early recovery and model recovery projections (Bozec et al. 2022) under climate change (Lachs et al. 2024)

### 1. Hydrodynamics influences coral settlement

The spatiotemporal patterns of larval supply remain largely uncharacterized on coral reefs, as only few studies have attempted to quantify the supply of coral larvae during their planktonic stage empirically (Black 1988, Oliver et al. 1992, Gilmour et al. 2009). Instead, research has predominantly focused on settlement to artificial substrates as a proxy for larval supply, thereby limiting our understanding of critical pre-settlement processes for corals (Fig. 1). This study did not quantify natural coral larval fluxes but instead manipulated the retention timeframe of larval supply by delivering larvae under different retention timeframes, the shortest representing the average slack current conditions found in coral reef habitats (Gouezo et al. 2025). At the high concentrations of competent larvae (22,000-44,000 larvae m^-2^) utilized in this study, the residence time over a reef during the ∼2-3 hour slack current period resulted in an elevated settlement signal. This finding suggests that the likelihood of transition from planktonic larval to sessile benthic phases are potentially highest during periods of slack currents for most reefs. These conditions create a temporal window of opportunity for competent coral larvae to swim downwards to the reef and become entrained within the rough reef matrix of the benthic boundary layer where current speed is minimal (Boudreau and Jorgensen 2001).

When current velocity in the water column exceeds a certain threshold, the slow swimming speed of coral larvae (Hata et al. 2017) may limit their ability to swim down to the benthos directly below. Coral larvae swim at speeds of 0.1 to 0.5 cm.s-1, which is very slow compared to most other marine invertebrate larvae (Chia et al. 1984). Larvae could experience horizontal displacement 100s meters downstream when swimming downward, depending on prevailing current speed and their distance from the benthos (Takeda-Sakazume et al. 2022)(Table S14). When current speed exceeds 0.06 m.s^-1^, and depending on reef size, larvae may fail to reach the targeted reef. Instead, they might continue drifting until encountering another reef, areas with slower water movement, or no substrate at all. This occurrence is influenced by the relationship between larvae swimming speed, reef size, horizontal current velocity (Table S14), and local turbulence and hydrodynamics (Fig. 1).

Our observation of a significantly elevated settlement signal following larval release during a 2–3-hour retention timeframe reveals a previously undocumented role of hydrodynamics in initiating coral recovery. In an ecological context, current speed conditions after the arrival of a larval plume drive larvae residence times near the reef and boost the likelihood of settlement if suitable substrates are available (Gouezo et al. 2025). This change in settlement likelihood could be better integrated into dispersal modelling studies. Most studies investigating metapopulation connectivity at reefal scales (Boschetti et al. 2020) consider the total number of larvae arriving at a reef as representative of settlement, failing to account for temporal changes in settlement probability with changing current speed. Therefore, incorporating probabilities of larval settlement that increase when water residence time at a reef extends (Greenwood et al. Under Review) would better reflect empirical measurements.

### 2. High larval retention promotes gregarious settlement, chimerism, and space limitation on tiles

A settlement signal was observed across all larval retention treatments, but was 4-7 times higher under the 2.5-and 24-hour retention treatments, leading to saturation of larvae within the 2.25 m^2^ study plots and extremely high densities of settlers on tiles (20 tiles showing 50- 158 settlers per tile). Settlement tiles induce may coral larval settlement (Harriott and Fisk 1987) due to crustose coralline algae and other inducers (Harrington et al. 2004, Price 2010) and tile surface microstructure providing refugia (e.g. Edmunds et al. 2014). Density-dependent settlement influences early coral survival both positively and negatively. Some corals exhibit gregarious settlement behaviour, leading to polyps aggregating into large colonies due to space limitation (Doropoulos et al. 2017, Sampayo et al. 2020). Gregarious settlement can result in fusion into chimeras in single-species hard and soft corals (Rinkevich 2004, Puill-Stephan et al. 2012). Our study demonstrates that chimerism also occurs in mixed-species assemblages dominated by Acroporidae. High chimeric formation rates in multi-species larval assemblages suggest larval allorecognition (Barki et al. 2002) or early-stage fusion, creating a ‘mosaic genotype’ and intra-colonial genetic variability (Oury and Magalon 2024). Chimeric formations led to faster growth and survival rates, consistent with previous studies using single species (Raymundo and Maypa 2004, Cameron and Harrison 2020, Harrison et al. 2021). Though not statistically significant (Fig. S6, S7), high settler density per tile may have contributed to negative density-dependent population regulation, as recruit loss was highest on densely populated tiles (Fig. 2, 3). Previous studies have reported similar patterns where high initial settlement densities led to lower recruit densities over time (Doropoulos et al. 2017, Cameron and Harrison 2020).

### 3. Microhabitat filtering and recruitment rates

The positive influence of chimerism observed on tiles during the initial 255 days appeared to have been counteracted by the tile microhabitat’s light conditions (Fig. 3 & 4). Regardless of larval retention treatments and initial settler densities, recruitment rates on the tiles from all treatments were similar after 455 days, averaging 0.5-1.3 recruits per tile. Coral recruits near the underside tile edge showed high survivorship, whether chimeric or not. All recruits surviving at 455 days were on the underside outer edge of the tile (∼1 cm band), growing over the edges, similar to Cameron and Harrison (2020). This suggests a potential microhabitat filtering effect within the tile environment (Fig. 4). As light dims toward the tile’s centre, it may select for coral recruits favouring specific light intensity and spectral quality (Mundy and Babcock 1998). Also, the central more-shaded conditions on the tile’s underside could facilitate colonial organisms like bryozoans or ascidians, which outcompete coral recruits in more light-limited zones (Fig. 4) (Maida et al. 1994, Doropoulos et al. 2016).

This study provides coral recruitment data on natural substrata using macrophotogrammetry (Gouezo et al. 2023) from 66 days after larval supply for the two longest and most concentrated larval retention treatments (2.5h and 24h net, 44,000 larvae per m^2^). No significant difference was observed in coral recruit densities.m^-2^ between treatments on natural substrates or tiles (Fig. 3, Table S7). Densities on the reef were 11-times lower than on tiles at 66 days post-settlement but only 3-times lower after 455 days. This suggests the high proportional loss of recruits on tiles was not evident on reef substrata (Fig. 4), which showed higher post-settlement survivorship and density-independent survival (Fig. S5, S6). Recruit counts on natural substrata were not statistically different between 255 and 455 days, whereas they significantly decreased on tiles.

At 15 months post-larval supply, recruitment on the reef reached an average of 47 recruits m^-2^. This value represents some of the highest reported *Acropora* densities on the substrata following coral larval enhancement (dela Cruz and Harrison 2017, 2020, Harrison et al.

2021), which are likely attributable to higher detectability using macrophotogrammetry during the early post-settlement phase (Gouezo et al. *in prep*). Interestingly, early-stage colony sizes of *Acropora* were lower than those reported in the Philippines for *Acropora tenuis* and *A. loripes* (dela Cruz and Harrison 2017, 2020, Harrison et al. 2021). At 13-15 months post-supply, recruit maximum diameter reached approximately 25-40 mm in the Philippines (dela Cruz and Harrison 2017, 2020, Harrison et al. 2021), while averaging about 7 mm (reef) and 10 mm (tile) on the Lizard Island fore reef (Fig. S8). This is potentially due to reduced chimerism with mixed species larvae assemblages, lower species-specific growth rates, and/or cooler seawater temperatures at Lizard Island than in northern Luzon, Philippines. While this study provides estimates of early-stage coral recruitment and growth rates on natural substrata, recruitment and growth varied considerably among reef plots (Fig. 3). Future work is needed to identify biophysical factors on natural substrata driving this variability to improve characterisations of the optimal recruitment niche.

In conclusion, our study shows that episodic slack current conditions lasting less than 3-hours sufficiently retained coral larvae locally, resulting in higher settlement of supplied cultured larvae compared to background rates. Despite initial differences in settlement rates across larval retention treatments, the number of surviving recruits after 15-months on tiles was similar, indicating post-settlement processes played a crucial role in regulating coral population densities. Coral recruitment on the reef exhibited 2.5- to 3.5-times higher survival than on tiles, suggesting that drivers of post-settlement bottlenecks were stronger on tiles than on reef substrata. These outcomes also indicate that tile-based assessments do not accurately reflect recruitment dynamics on the reef. While our study provides valuable insights, further research is necessary to confirm whether these patterns are representative across diverse coral reef habitats and ecosystems. Our findings underscore the need to investigate the influence of pre-versus post-settlement processes on recruitment success to elucidate how they modulate each other in driving early coral recovery and improve modelling of reef recovery trajectories, which currently lack empirical data on supply-settlement relationships across different reef habitats.

## ACKNOWLEDGEMENTS

We acknowledge the Traditional Owners of Lizard Island on which this research was conducted, the Dingaal, Thaanil-Warra and Ngurruumungu Aboriginal people. We thank staff from SCU, CSIRO, Blue Planet Marine and Lizard Island Research Station for providing support during field work activities. This research was funded by the Moving Corals subprogram of the Reef Restoration and Adaptation Program (RRAP) to PH and CD, and the grant ERRFP-1198 awarded to George Roff. RRAP is funded by the partnership between the Australian Government’s Reef Trust and the Great Barrier Reef Foundation. This study was conducted under GBRMPA permit no. G20/44511.1.

## AUTHOR CONTRIBUTIONS

MG, PH, GR, and CD conceived the study idea and designed methodology; MG, AC, CD, DT, LH, and AF conducted the field experiment and collected data; MG, BG, MG, and GR developed the tile imagery analysis workflow; MG and AC extracted data from imagery, MG conducted all analyses and led the draft of the manuscript. CD, GR, DT and PH contributed critically to the manuscript drafts. All authors gave final approval for publication

## CONFLICT OF INTEREST STATEMENT

The authors declare no conflict of interest.

## Notes

### Competing Interest Statement

The authors have declared no competing interest.

