## Supplementary Information for "The influence of larval retention on coral recruitment"

**Supplementary Figures**

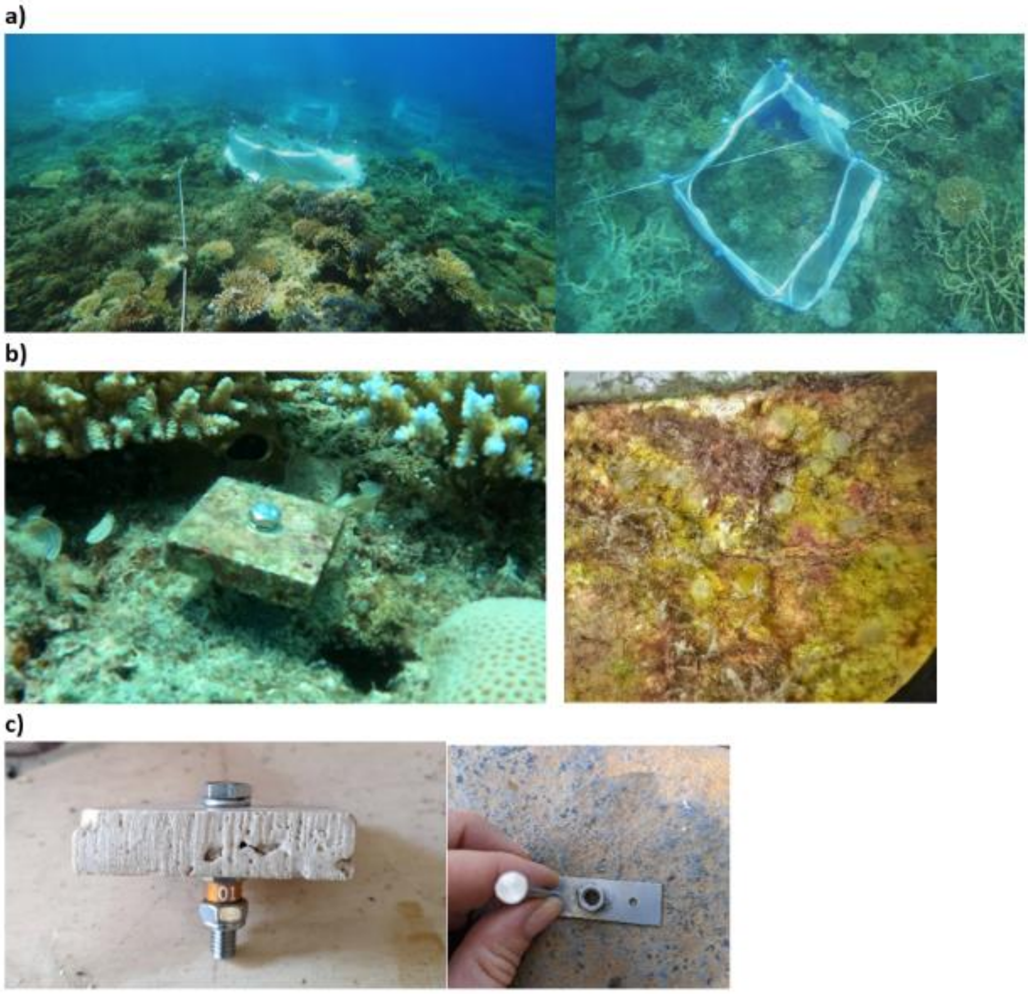

**Figure S1: (a) 1.5 m x 1.5 m experimental plots randomly located on either side of a 50 m transect set at 5 m depth, showing when the net was opened after 2.5 h of retention. (b) Example settlement tile secured ~2 cm off the substrate on a baseplate, with a high-resolution image of 2-day old settlers following supply treatments. (c) The set-up of the tile deployment system, assembled using M6 stainless bolts, washers, lock nuts, and tags, and deployed onto the substrate into small stainless-steel baseplates with an M6 nut welded in the centre were nailed to the substrate using 2.8 mm galvanized nails.**

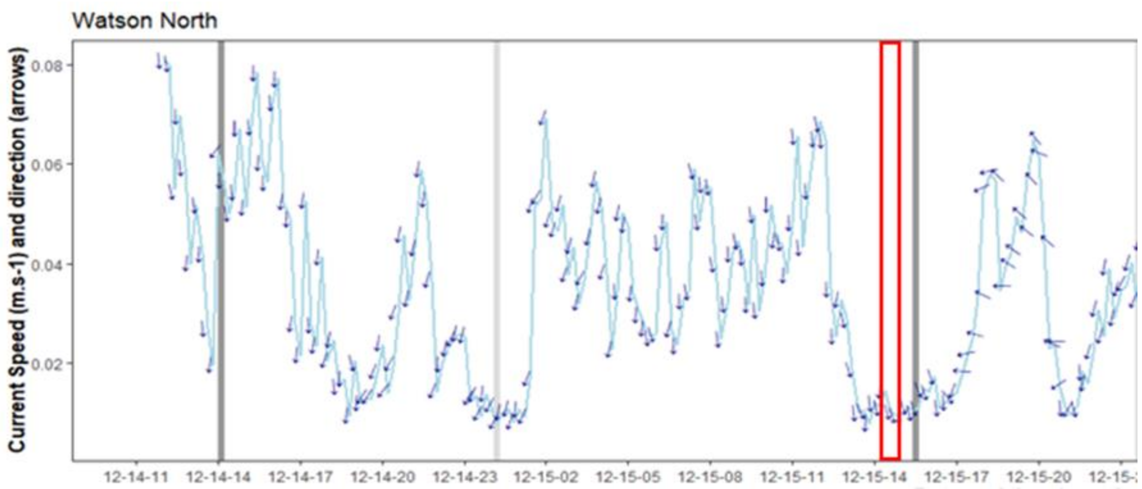

**Figure S2. Current condition at experimental site prior, during (red square) and following larval supply treatments. Dark grey bar shows the timing of high tide and light grey bar shows the timing of low tide.**

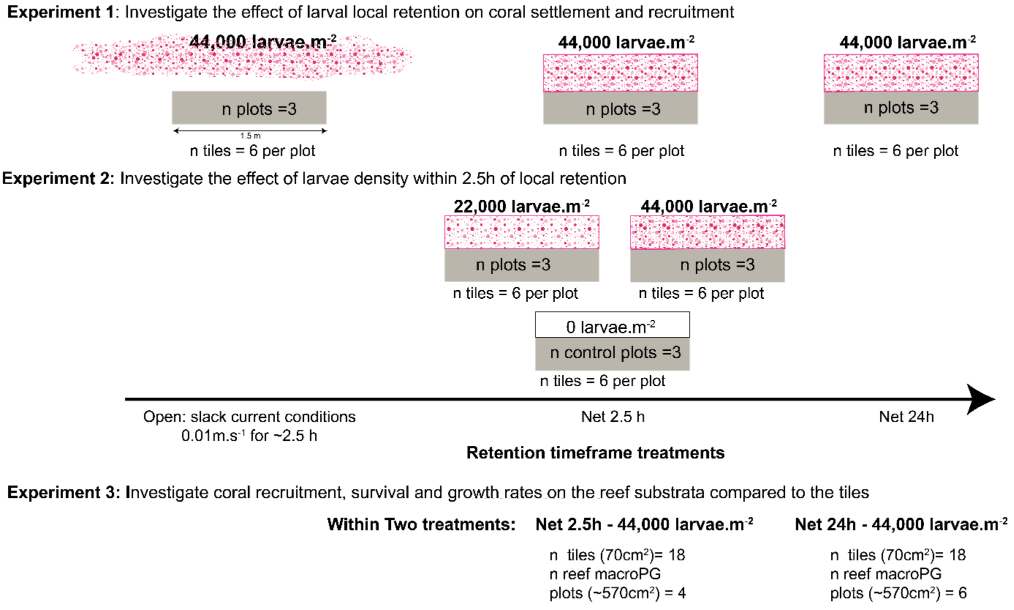

**Figure S3: Diagram showing the overall experimental design, describing the purpose of each of the three experiments. Current conditions on the reef on the day of the experiment are shown in Figure S1.**

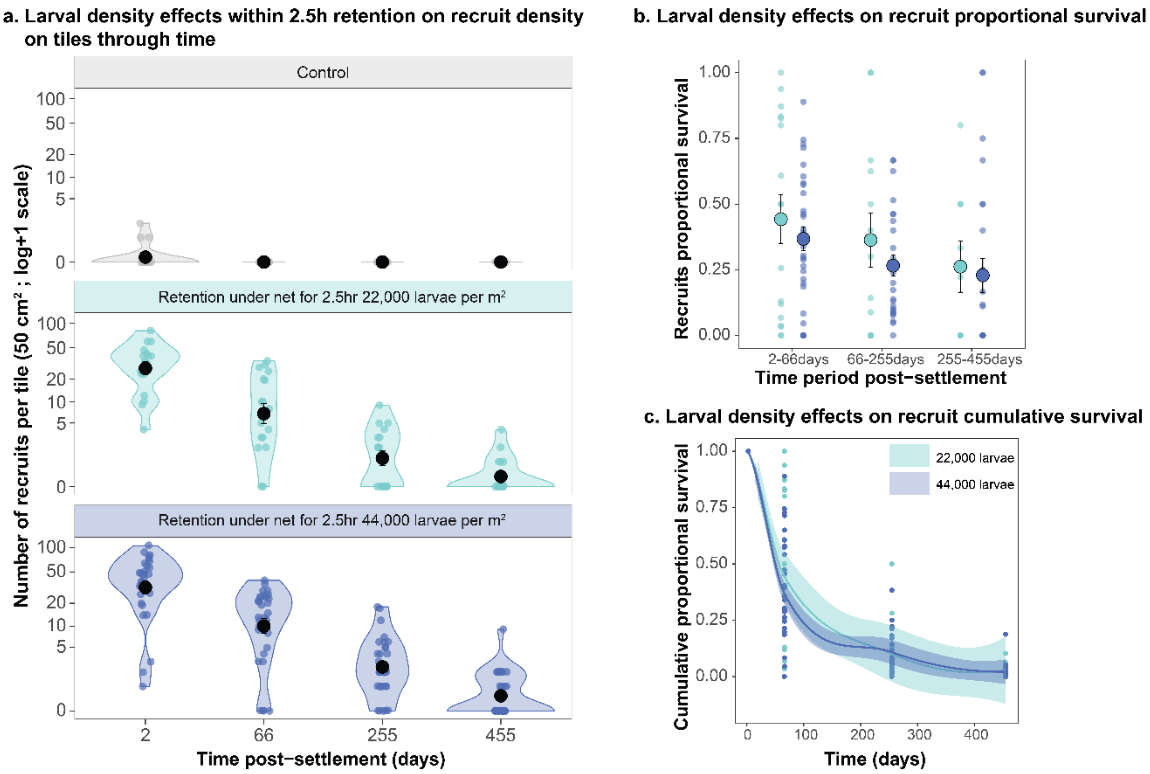

**Fig S4**: **Larval density within 2.5h retention treatments effect on coral recruit (a) densities, (b) proportional survival, and (c) cumulative survival on tiles.**

a. Time series and photographs of coral recruits on reef substrate

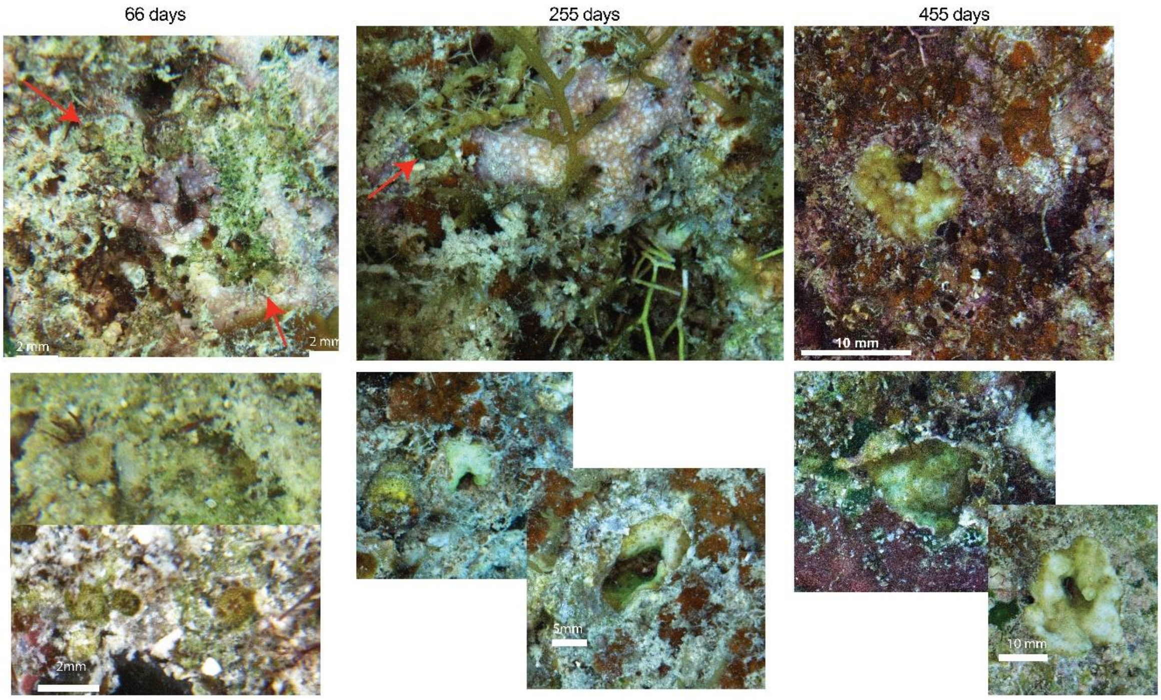

b. Time series and photographs of coral recruits on the underside face of the tile

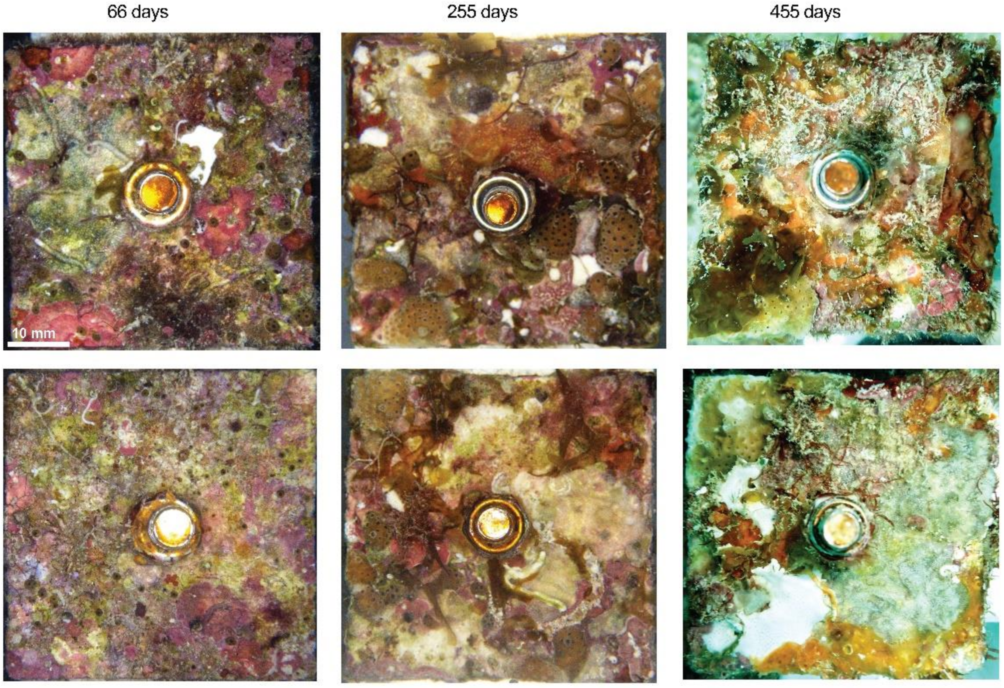

**Fig S5. Photographs of the reef (a) and the underside of tiles (b) showing coral recruits at the three monitoring time points**

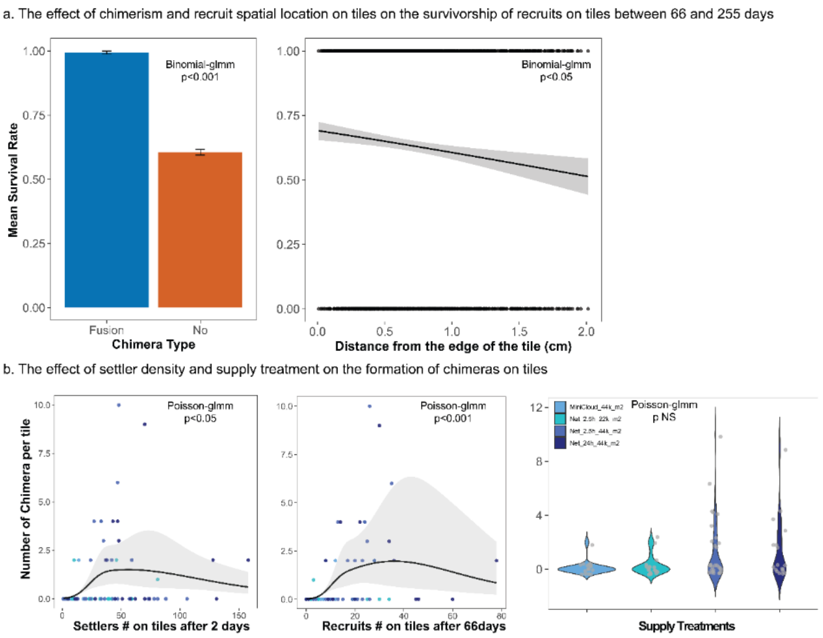

**Figure S6: Plots showing the significant findings from the analysis investigating the effect of chimerism, density dependence, and recruit location within the tile grid on the survivorship of recruits on tiles (a), and the effect of settler densities at both 2 and 66 days on tiles and supply treatment on the number of chimeras (b)**

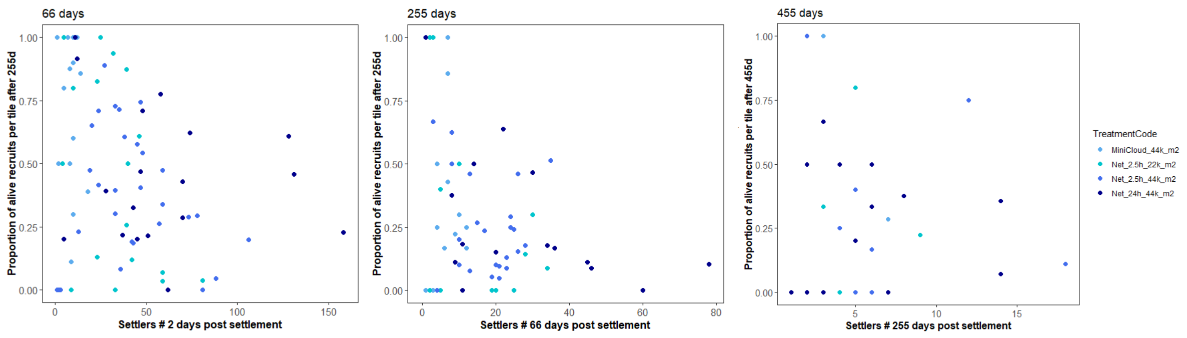

**Figure S7. The effect of recruit density on the proportion of surviving recruits for each monitored time period: 2-66d ; 66-255d ; 255-455 day (All relationships non-significant (Table S.11, S12)**

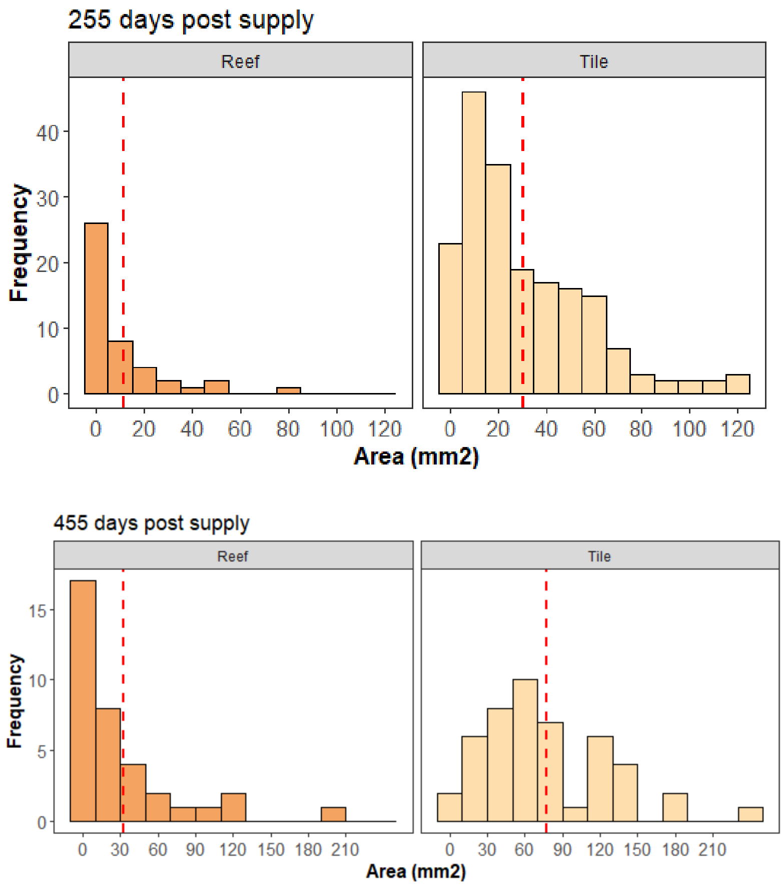

**Figure S8. Size distribution of coral recruits following two larvae supply treatments. Diameter inferred using the formula: $2r = 2\sqrt{(A / \pi)}$, average diameter of 6.5 and 9.9 mm for reef and tile respectively after 455 days post settlement.**

**Supplementary Tables**

### Objective 1: Larval retention effects on coral recruitment

Table S1: Effect of retention treatments on initial settler density on tiles

| RecID_count ~ TreatmentCode + (1 \| PlotID),<br>family = nbinom1()) | | | | |
|---|---|---|---|---|
| | **statistic** | **df** | **p.value** | contrasts |
| Retention Treatments | 54.79034 | 3 | <0.001 | Control < Cloud < Net_2.5h = Net_24h |

Table S2: Effect of retention and treatments on recruitment on tiles through time

| Rec_count ~ TreatmentCode*Time + (1 \| PlotID),<br>family = nbinom2 | | | | |
|---|---|---|---|---|
| | Chisq | Df | P-values | contrasts |
| Retention Treatments | 7.065 | 2 | 0.029 | Open < Net_24h |
| Time | 220.130 | 2 | <0.001 | 66 > 255 > 455 d |
| Retention Treatments:Time | 3.676 | 4 | 0.452 | |

Table S3: Effect of retention treatments on recruit survivorship on tiles through time

| status ~ TreatmentCode + (1 \| PlotID/TileRep), family = binomial() | | | | | |
|---|---|---|---|---|---|
| Time period | Fixed effect | statistic | df | p.value | contrasts |
| 2-66 | Retention Treatments | 10.639 | 2 | 0.00489345 | Open > Net Treatments |
| 66-255 | Retention Treatments | 4.24133868 | 2 | 0.119951313 | NS |
| 255-455 | Retention Treatments | 0.102002509 | 2 | 0.950277478 | NS |

**Objective 2: Larval density effects on coral recruitment**

Table S4: Effect of larval density within 2.5-h retention treatments on initial settler density on tiles

| Rec_count ~ TreatmentCode + (1 \| PlotID), family = nbinom1() | | | | |
|---|---|---|---|---|
| | statistic | df | p.value | contrasts |
| Density Treatment within 2.5h retention | 36.66495635 | 2 | 1.09221E-08 | Control < Net_2.5h_22K = Net_2.5h_44K |

Table S5: Effect of larval density within 2.5-h retention treatments on recruit densities on tiles through time

| Rec_count ~ TreatmentCode*Time + (1 \| PlotID), family = nbinom2 | | | | |
|---|---|---|---|---|
| | Chisq | Df | P-values | contrasts |
| Treatment | 0.824 | 1 | 0.364 | |
| Time | 131.894 | 2 | <0.001 | 66 > 255 > 455 d |
| Treatment:Time | 0.426 | 2 | 0.808 | |

Table S6: Effect of larval density within 2.5-h retention treatments on recruit survivorship on tiles through time

| status ~ Treatment + (1 \| PlotID/TileRep), family = binomial() | | | | | |
|---|---|---|---|---|---|
| Time period | Fixed effect | statistic | df | p.value | contrasts |
| 2-66 | Treatment | 0.651759432 | 1 | 0.419484338 | |
| 66-255 | Treatment | 0.113380603 | 1 | 0.736327711 | |
| 255-455 | Treatment | 0.03019896 | 1 | 0.862039559 | |

**Objective 3: Substrate effects on coral recruitment**

Table S7: Effect of the two highest retention treatments on recruit counts on the substrata

| Rec_count ~ TreatmentCode*Time + offset(SampleArea_m2), family = gaussian | | | | |
|---|---|---|---|---|
| term | statistic | df | p.value | |
| Treatment | 0.684 | 1 | 0.408 | |
| Time | 13.738 | 2 | 0.001 | 66 d = 255 d < 455d (marg)<br>66 d < 455 d |
| Treatment:Time | 0.410 | 2 | 0.815 | |

Table S8: Effect of the substrate type on the count of recruits through time

| Rec_count ~substrate type *Time + offset(log(SampleArea_m2) + (1\|PlotID), family = nbinom2() | | | | |
|---|---|---|---|---|
| term | statistic | df | p.value | contrasts |
| Substrate type | 144.072 | 1 | 3.43E-33 | |
| Time | 283.3155 | 2 | 3.01E-62 | |
| Substrate type: Time | 10.58009 | 2 | 0.005042 | Within Time: All, Reef < Tile<br>Within Substrate type<br>For reef, 66 < 255=455<br>For Tile, 66 < 255 < 455 |

Table S9. Effect of recruitment substrate type on recruit's survivorship through time

| Status ~ substrate type + (1\| PlotID), family = binomial("logit") | | | | | |
|---|---|---|---|---|---|
| Time period | Fixed effect | **statistic** | **df** | **p.value** | |
| 66-255 days | substrate type | 40.64848375 | 1 | 1.82229E-10 | Tile< MP |
| 255-455 days | substrate type | 22.73638081 | 1 | 1.85816E-06 | Tile< MP |

Table S10. Effect of recruitment habitat and fusion on recruit's growth rate

| Growth rate ~ substrate type _Chimera + (1 \| PlotID), family = Gamma(link= "log") | | | | | |
|---|---|---|---|---|---|
| Time period | Fixed effect | **statistic** | **df** | **p.value** | contrasts |
| 66-255 days | substrate type _Chimera | 22.11719 | 2 | 1.58E-05 | Reef_no Fusion < Tile_no fusion < Tile_ fusion |
| 255-455 days | substrate type _Chimera | 3.648158871 | 2 | 0.161366124 | NS |

**Objective 4: Tile-specific drivers of mortality (Objective 4)**

Table S11. Recruit survivorship on tile habitat

<table>
<tr><td colspan="5">Surv ~ Rec_count_202212+(1|TileRep)+(1|PlotID),<br>family = binomial("logit"))</td></tr>
<tr><td>Time period</td><td>Fixed effect</td><td>statistic</td><td>df</td><td>p.value</td></tr>
<tr><td>0-66 days</td><td>Rec_count_202212</td><td>0.565</td><td>1</td><td>0.452</td></tr>
</table>

Table S12. Recruit survivorship on tile habitat

<table>
<tr><td colspan="7">Surv ~ Chimera+distance_from_edge+Rec_count+(1|TileRep)+(1|PlotID),<br>family = binomial("logit"))</td></tr>
<tr><td>Time period</td><td>Fixed effect</td><td>statistic</td><td>df</td><td>p.value</td><td>contrasts</td><td>R2</td></tr>
<tr><td>66-255 days</td><td>Chimera</td><td>20.4929</td><td>1</td><td>5.99E-06</td><td>No<Yes</td><td rowspan="3">0.31</td></tr>
<tr><td></td><td>distance_from_edge</td><td>6.10972</td><td>1</td><td>0.013444</td><td></td></tr>
<tr><td></td><td>Rec_count_202302</td><td>0.0175</td><td>1</td><td>0.89</td><td>NA</td></tr>
<tr><td>255-455 days</td><td>Chimera</td><td>3.86</td><td>1</td><td>0.0494</td><td>No<Yes</td><td rowspan="3">0.01</td></tr>
<tr><td></td><td>distance_from_edge</td><td>0.0140</td><td>1</td><td>0.906</td><td>NA</td></tr>
<tr><td></td><td>Rec_count_202308</td><td>1.75</td><td>1</td><td>0.186</td><td>NA</td></tr>
</table>

Table S13. Chimera counts as a function of initial density and treatments

<table>
<tr><td colspan="6">Chimera ~ ns(RecID_count_202212, df = 3)+Treatment,<br>ziformula = ~1,<br>family = poisson())</td></tr>
<tr><td>Fixed effect</td><td>statistic</td><td>df</td><td>p.value</td><td>contrasts</td><td>R2</td></tr>
<tr><td>ns(Rec count_202212, df = 3)</td><td>8.531997</td><td>3</td><td>0.036206</td><td>~peak @ 50</td><td rowspan="2">0.88</td></tr>
<tr><td>Treatment</td><td>4.467898</td><td>3</td><td>0.215172</td><td>NA</td></tr>
<tr><td>ns(Rec count_202302)</td><td>13.13008139</td><td>3</td><td>0.004363577</td><td>~peak @ 25</td><td>0.94</td></tr>
<tr><td>Supply treatments</td><td>2.684956102</td><td>3</td><td>0.442789904</td><td></td><td></td></tr>
</table>

Table S14. shows a simplified estimation of horizontal displacement of larvae when larvae decide to swim to the reef given their location in the water column and their swimming speed. This does not take into account local turbulences near the reef benthos that may influence the pace at the which the reef benthos is encountered.

| Vertical distance between larvae and reef (m) | Time (s) to swim downward if larvae swimming speed is ~ 2mm/s | Horizontal displacement (m) during slack current = 0.01 m/s | Horizontal displacement (m) for highest current speed = 0.07 m/s |
|---|---|---|---|

| | | | |
|---|---|---|---|
| 1 | 500 s (8.3 min) | 5 | 35 |
| 2 | 1000 s (16.7 min) | 10 | 70 |
| 3 | 1500 s (25 min) | 15 | 105 |
| 4 | 2000 s (33 min) | 20 | 140 |
